# Human primosome requires replication protein A when copying DNA with inverted repeats

**DOI:** 10.1101/2024.03.11.584335

**Authors:** Andrey G. Baranovskiy, Lucia M. Morstadt, Eduardo E. Romero, Nigar D. Babayeva, Tahir H. Tahirov

**Affiliations:** Eppley Institute for Research in Cancer and Allied Diseases, Fred & Pamela Buffett Cancer Center. University of Nebraska Medical Center, Omaha, NE, USA; Department of Biochemistry, University of Nebraska-Lincoln, Lincoln, NE, USA; Nebraska Center for Biotechnology, University of Nebraska-Lincoln, Lincoln, NE, USA; Nebraska Center for Biomolecular Communication (CIBC), University of Nebraska-Lincoln, Lincoln, NE, USA

**Keywords:** Primosome, DNA polymerase α, RPA, winged helix-turn-helix, DNA hairpin, inverted repeats, G-quadruplex, DNA polymerase ε, protein-DNA interaction, protein-protein interaction, binding studies, cryo-EM, structure

## Abstract

The human primosome, a four-subunit complex of primase and DNA polymerase alpha (Polα), initiates DNA synthesis on both chromosome strands by generating chimeric RNA-DNA primers for loading DNA polymerases delta and epsilon (Polε). Replication protein A (RPA) tightly binds to single-stranded DNA strands, protecting them from nucleolytic digestion and unauthorized transactions. We report here that RPA plays a critical role for the human primosome during DNA synthesis across inverted repeats prone to hairpin formation. On other alternatively structured DNA, forming a G-quadruplex, RPA does not assist primosome. A stimulatory effect of RPA on DNA synthesis across hairpins was also observed for the catalytic domain of Polα but not of Polε. The winged helix-turn-helix domain of RPA is essential for an efficient hairpin bypass and increases RPA-Polα cooperativity on the primed DNA template. Cryo-EM studies revealed that this domain is mainly responsible for the interaction between RPA and Polα. The flexible mode of RPA-Polα interaction during DNA synthesis implies the mechanism of template handover between them when the hairpin formation should be avoided. This work provides insight into a cooperative action of RPA and primosome on DNA, which is critical for DNA synthesis across inverted repeats.

**SIGNIFICANCE:** This work revealed the critical role of RPA during DNA synthesis across inverted repeats by the human primosome. It was shown that a small winged helix-turn-helix domain of RPA is essential for the cooperative action of RPA and primosome, especially when copying DNA sequences prone to hairpin structure formation. Structural studies uncovered the mode of RPA and Polα integration, which assures an efficient DNA handover between them. This work provides a notable insight into the early steps of Okazaki fragments synthesis.

## INTRODUCTION

The human primosome is a multifunctional complex that plays a key role in genome replication (1, 2). It initiates DNA synthesis at every origin of replication by generating chimeric RNA-DNA primers essential for recruiting two main replicases: DNA polymerase epsilon (Polε) and delta (Polδ). Primosome is involved in a variety of other cellular processes, including telomere maintenance (3), genome stability (4, 5), and innate immunity (6). It is a promising candidate for antitumor therapy (7).

The human primosome is a complex of two enzymes: the two-subunit primase, with catalytic (p49) and regulatory (p58) subunits, and the two-subunit DNA polymerase α (Polα), with catalytic (p180) and accessory (p70) subunits (1, 2). p180 contains a catalytic domain (Polα_CD_) and a C-terminal domain (p180_C_); p58 has N-terminal (p58_N_) and C-terminal (p58_C_) domains. Primosome forms an elongated platform p49-p58_N_-p180_C_-p70, which interacts with a CMG helicase (8) and holds p58_C_ and Polα_CD_ by linkers with a length of ∼20 amino acids each (9, 10). Polα_CD_ adopts the universal “right-hand” DNA polymerase fold with an active site formed by a “palm” holding the catalytic aspartates, a “thumb” that binds the template:primer (T:P) duplex, and “fingers” interacting with an incoming nucleotide (11).

The primase is responsible for the initiation, elongation, and termination steps of RNA synthesis (1, 12, 13). We have shown that human primase is structurally predetermined to synthesize an RNA primer with a length of nine nucleotides (9). Owing to very low affinity of p49 to the primer 3’-end (14, 15), Polα takes over the 9-mer RNA primer and extends it by ∼25 deoxyribonucleotides (16, 17). Importantly, p58_C_ tightly binds to the primer 5’-end (15, 17) and is a major regulator of all primosome transactions, also serving as a processivity factor for primase and Polα (9, 10, 18, 19).

Replication protein A (RPA) is a hetero-trimeric protein with a conserved domain organization. It is a major single-stranded DNA (ssDNA) binding protein in eukaryotes. RPA protects unwound DNA strands from nucleolytic damage and prevents reannealing and the formation of alternate structures (20, 21). RPA participates in various DNA metabolic processes involving ssDNA, including DNA replication, recombination, and repair, telomere maintenance, DNA damage response, and checkpoint activation (22). A recent study utilizing reconstituted reactions with yeast proteins has shown that RPA is critical for origin unwinding and DNA replication (23).

Human RPA is composed of three subunits: RPA70, RPA32, and RPA14 (Fig. 1), and contains six oligonucleotide-binding domains (F, A, B, C, D, and E), with four of them (A-D) being involved in ssDNA-binding (21). RPA comprises two main parts that are flexibly connected: N-terminal (domains F, A, and B) and C-terminal (RPAcore; contains both small subunits and the C-domain of RPA70). A zinc-binding motif is a part of the C-domain and is important for structural stability and DNA binding (24, 25). Besides DNA binding, C and D domains also participate in intersubunit interactions, forming with RPA14 a minimal trimerization core (RPAcore^ΔP,W^) (26). The disordered N-terminus of RPA32 (P-motif; residues 1-33) undergoes phosphorylation in response to cell cycle progression and DNA damaging agents, however, the structural and functional impact of this modification is poorly understood (27). The C-terminus of RPA32 contains a flexibly tethered winged helix-turn-helix domain (wHTH- or W-domain; residues 203-270), which interacts with different DNA processing proteins with relatively low affinity (*K_D_* in the range of 5-10 µM) (28, 29). The N-terminal part of RPA70 has several binding partners, including viral proteins (21).

**Figure 1.**
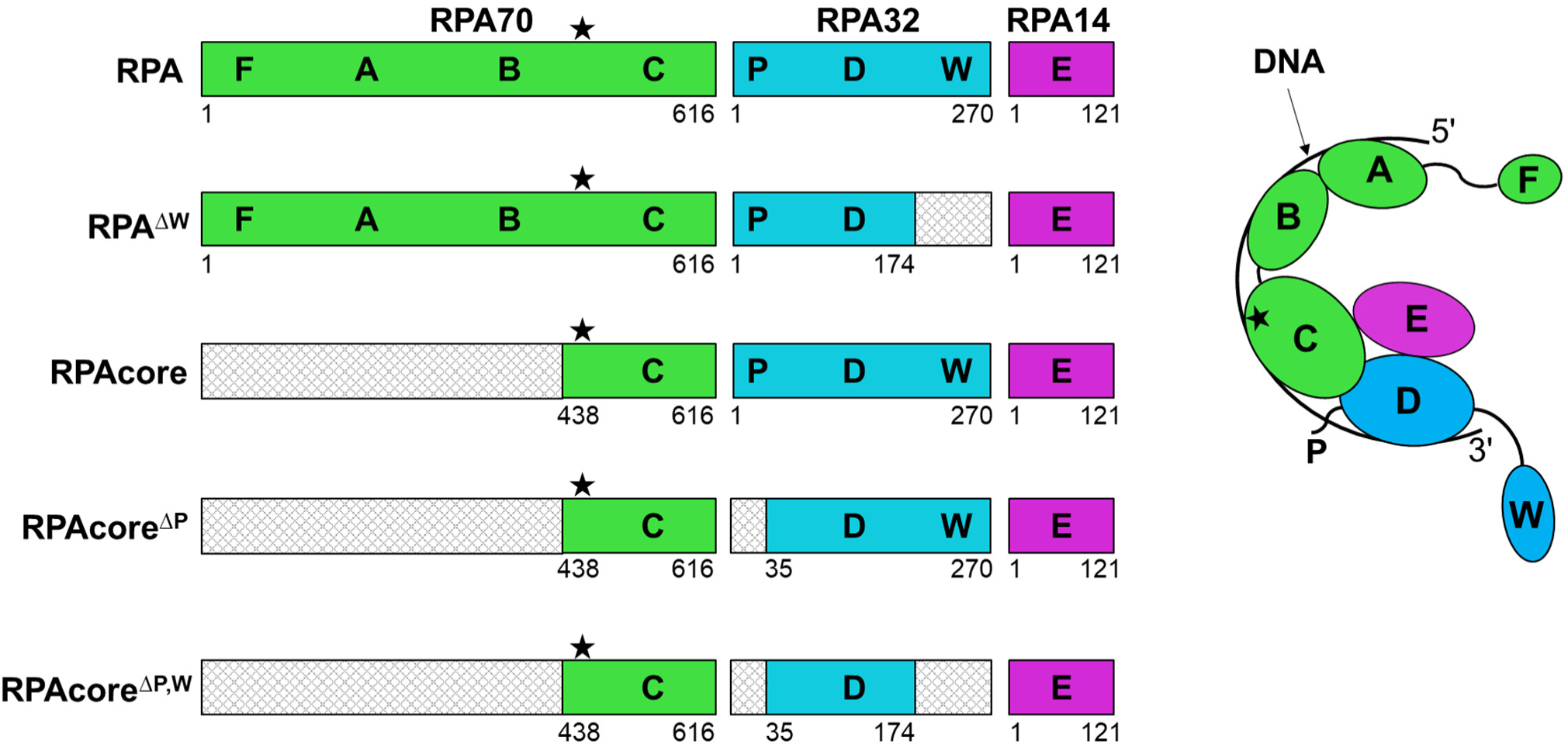
A schematic representation of the domain organization of human RPA and its variants and the mode of RPA-DNA interaction. RPA contains six domains (F, A, B, C, D, E) and four (A-D) participate in DNA binding. RPAcore is a complex of two small subunits and the C-domain of RPA70. The star symbol denotes a zinc-binding motif, P – phosphorylation motif, and W – winged helix-turn-helix domain (wHTH). The deleted domains are depicted by a cross-hatching pattern.

RPA binds ssDNA with high affinity, with *K_D_* values in the sub-nanomolar range (30). According to structural data, the AB-domains and RPAcore interact with 8 and 17 nucleotides, respectively (31, 32). RPA binds ssDNA with a defined 5’→3’ polarity, with small subunits facing the T:P junction (33). Of note, RPAcore retains the ability to recognize DNA polarity (34). An interaction between RPA and primosome was detected over thirty years ago using enzyme-linked immunoassay (35). Later, a more elaborated study has shown that the W-domain plays an important role in RPA-Polα interaction (36).

This work shows that RPA strongly stimulates primosome and Polα_CD_ upon DNA synthesis across inverted repeats. We conducted biochemical experiments and cryo-EM studies to gain insight into the mechanism of RPA-Polα cooperation and a DNA template handover between them. Several RPA mutants were generated to study the role of RPA domains on the activity of Polα alone and in complex with primase, in regard to the structure of a DNA template. Binding studies were conducted to explore the W-domain’s role in stabilizing of the RPA/Polα/DNA complex. Altogether, our studies revealed the mode of RPA-Polα interaction and the importance of their cooperation during DNA synthesis.

## RESULTS

### RPA inhibits primosome on unstructured DNA

We used a recently published assay utilizing a natural primer to analyze the effect of RPA and DNA secondary structures on primosome activity (17). In this assay, Polα extends the primer 3’-end while the primase accessory subunit holds the primer 5’-end. Primosome and RPA were pre-incubated with a 98-mer DNA template annealed to a 12-mer chimeric RNA-DNA primer containing the 5’-triphosphate (*SI Appendix*, Table S1), and DNA synthesis was initiated by addition of 10 µM dNTPs. The length of the 5’-tail is 81 nucleotides, which allows loading of at least two RPA molecules.

In primosome, the tight complex between the primase and the primer 5’-end allows Polα to perform processive DNA synthesis on the unstructured template (Fig. 2A, lanes 1 and 2). Simultaneous primer binding at the 5’-end by primase and at the 3’-end by Polα sets a limit for the length of a chimeric RNA-DNA primer defined by primosome linkers (10, 19). Of note, the primosome mainly generates 32-35-mer primers at single-turnover conditions (17).

**Figure 2.**
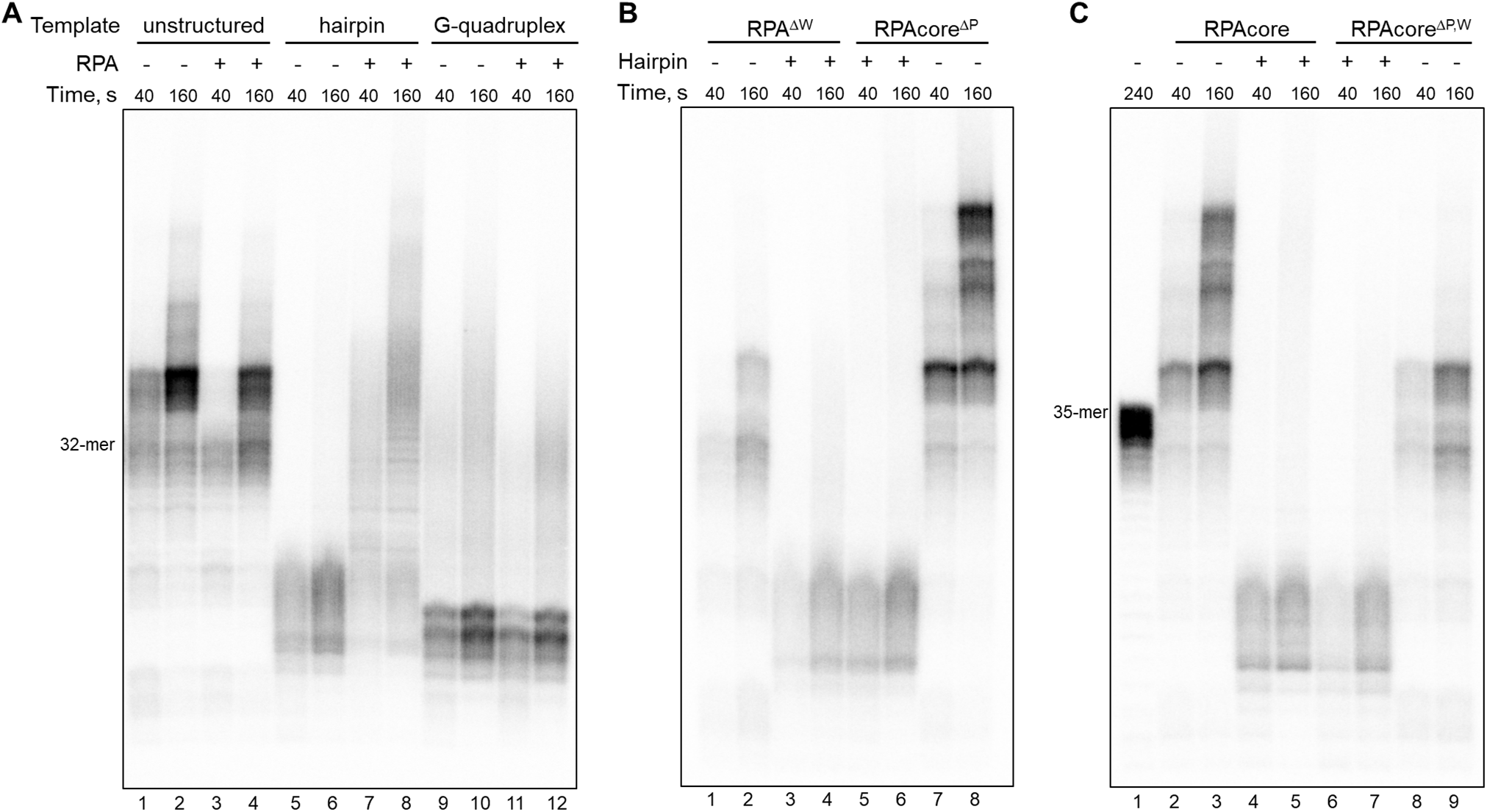
The role of RPA and its domains in DNA synthesis by primosome on DNA templates with and without secondary structures. **(A)** RPA stimulates the primosome upon DNA synthesis across a hairpin. **(B)** The W-domain and DNA-binding AB-domains are necessary for HP bypass by RPA/primosome. (**C)** RPAcore^ΔP,W^ does not stimulate primosome on both templates. The 40-mer template (T4) generated a 35-mer product (lane 1). In all other lanes, 98-mer DNA templates of different structures were annealed to a 12-mer chimeric RNA-DNA primer (P1) containing the triphosphate at the 5’-end. The following templates were used: T1 (unstructured or normal), T2 (hairpin), and T3 (G-quadruplex). Reactions were incubated at 35°C at specified time points.

On the unstructured (normal) template, RPA exhibits an inhibitory effect on DNA synthesis (Fig. 2A, lanes 1-4). Seemingly, RPA tightly bound to DNA should be an obstacle for DNA polymerases. Interestingly, RPA reduces the level of products with a length exceeding 32 nucleotides (Fig. 2A, lanes 1-4). This observation indicates that RPA prevents Polα from resuming DNA synthesis and is an additional factor in the composite mechanism of DNA synthesis termination by primosome (10, 17), which limits the amount of mutagenic DNA generated by Polα (4, 37).

### RPA helps primosome to bypass a hairpin but not a G-quadruplex

Next, we analyzed how RPA modulates primosome activity on alternatively structured DNA, which usually presents a challenge for DNA-processing proteins (38). In the template, structured DNA elements were placed nine nucleotides apart from the primer 3’-end to observe primosome stalling at the obstacle before DNA synthesis termination at the 32-mer product. A stable hairpin (HP) with a 9-bp stem completely stalls the primosome (Fig. 2A, lanes 5 and 6). Strikingly, RPA greatly stimulates HP bypass by the primosome (Fig. 2A, lanes 7 and 8).

The other known replication block tested was the G-quadruplex (G4) motif, which folds into a four-stranded structure composed of stacked guanine tetrads (39). G4 motifs are positioned non-randomly in the genome and are usually located at telomeres, replication origins, recombination sites, and gene promoter regions, including promoters of many oncogenes. The G4-forming motif stalls the primosome, and RPA shows no assistance (Fig. 2A, lanes 9-12). It is known that RPA can passively unfold hairpins and G4 structures by stabilizing the unfolded state (40–42). The failure of RPA to help Polα in DNA synthesis across a G4 motif can be attributed to the high stability of the G-quadruplex, resulting in a slow unfolding rate (40, 43).

### The W-domain is required for the hairpin bypass by primosome

Using an enzyme-linked immunosorbent assay, it was shown that the W-domain deletion disrupts the RPA-Polα interaction (36). The deleted region (residues 223-270) is a part of the W-domain (Fig. 1), which is flexibly connected to the rest of RPA and does not participate in the interaction with DNA. This domain mediates RPA interaction with different proteins, including the DNA recombination protein Rad52, the DNA repair proteins XPA and uracil DNA glycosylase 2, and the T-antigen of polyomavirus SV40 (21).

We generated an RPA variant named RPA^ΔW^ (Fig. 1 and *SI Appendix*, Fig. S1) by deleting the region 175-270, which contains the W-domain and the linker connecting it to the rest of RPA32. This deletion almost wholly abrogated the stimulatory effect of RPA on HP bypass by the primosome (Fig. 2B, lanes 3 and 4). These data show the critical role of the W-domain in DNA synthesis across HP by primosome. Of note, a W-domain deletion increased the inhibitory effect of RPA on unstructured DNA (Fig. 2B, lanes 1 and 2, in comparison to Fig. 2A, lanes 3 and 4, and *SI Appendix*, Fig. S2). Consistently, the recent study utilizing *in vitro* reconstituted DNA replication reactions with yeast proteins, showed a stimulatory role of the W-domain on the leading and lagging strand synthesis (23).

### RPAcore stimulates DNA synthesis but not a hairpin bypass

It is expected that the tight RPA-DNA interaction, which requires all four DNA-binding domains, is important for stabilizing unfolded hairpins and, therefore, for efficient HP bypass by the primosome/RPA complex. To test this idea, we generated the RPA variant RPAcore^ΔP^ by deleting the N-terminal part of RPA70 (residues 1-437; contains DNA-binding domains A and B), as well as the disordered N-terminus of RPA32 (P-motif; residues 1-34), whose function is not well characterized except that it becomes phosphorylated after DNA damage (27). The P-motif deletion significantly increased the protein yield after purification.

RPAcore^ΔP^ almost completely lost the ability to stimulate HP bypass (Fig. 2B, lanes 5 and 6). Interestingly, it shows the opposite effect on unstructured DNA versus the wild-type RPA (Fig. 2B, lanes 7 and 8, compared to Fig. 2A, lanes 3 and 4, and *SI Appendix*, Fig. S2), stimulating the DNA-polymerase activity. This fact supports the idea that high RPA affinity to DNA complicates its displacement by DNA polymerase.

We obtained two derivatives of RPAcore^ΔP^, one with intact RPA32 (named RPAcore) and the other with an additionally deleted W-domain (named RPAcore^ΔP,W^), which represents a minimal trimerization core comprises domains C, D, and E (26). RPAcore, like RPAcore^ΔP^, shows a stimulatory effect only on the unstructured DNA template (Fig. 2C, lanes 2-5). RPAcore^ΔP,W^, missing the W-domain, wholly lost the ability to stimulate primosome activity on both HP and a normal template (Fig. 2C, lanes 6-9). Thus, the stimulatory effect of RPA on HP bypass by primosome is dependent on the W-domain and the high affinity of RPA to DNA.

### The catalytic domain of Polα depends on the W-domain of RPA upon DNA synthesis across a hairpin

Next, we checked whether Polα_CD_ exhibits a similar dependence on RPA upon DNA synthesis across HP, as was observed for primosome. A previously published assay was employed where Polα_CD_ extends a 16-mer DNA primer annealed to a 73-mer template (44). On a normal template, Polα_CD_ generates in 1 minute a significant amount of a 70-mer primer (Fig. 3A, lane 2). In contrast to the primosome assay, DNA synthesis termination at the primer length of ∼35-nt is not observed due to the absence of primase and the triphosphate group on the primer (17). Polα_CD_ displays the same behavior on HP and G4 templates as primosome, making a pronounced pause at these structures (Fig. 3A, lanes 6, 7, 10, and 11). RPA does not stimulate Polα_CD_ on the normal template (Fig. 3A, lanes 2-5) but provides remarkable assistance for DNA synthesis across HP (lanes 6-9). As observed for primosome, RPA does not help Polα_CD_ on the G4 template (Fig. 3A, lanes 10-13).

**Figure 3.**
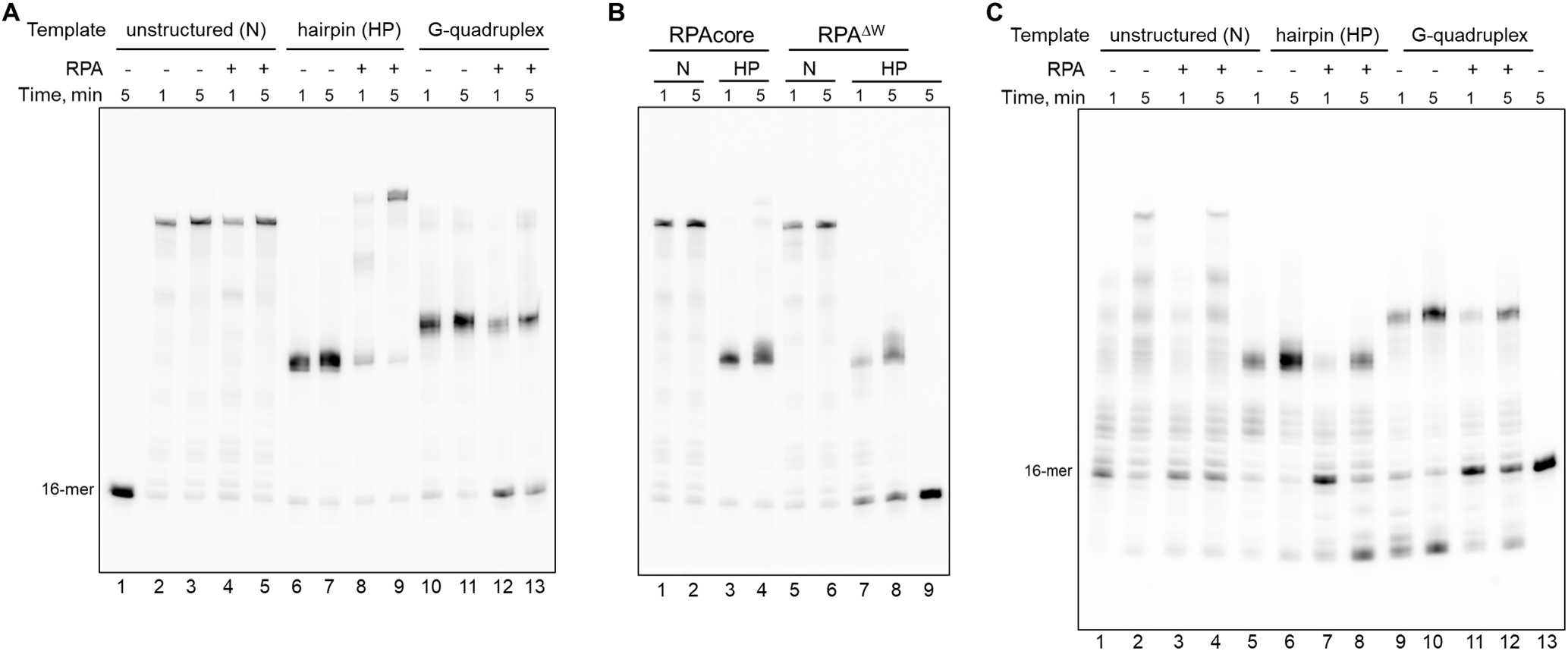
RPA assists Polα_CD_ and inhibits Polε_CD_ during DNA synthesis on the hairpin-containing template. (**A**) RPA stimulates Polα_CD_ on HP but inhibits it on the G4 template. (**B**) The W-domain and DNA-binding domains of RPA are critical for HP bypass by Polα_CD_. (**C**) RPA does not help Polε upon copying HP and G4 templates. DNA templates of different structures were annealed to a 16-mer DNA primer (P2) containing the Cy-5 fluorescent dye at the 5’-end. The following templates were used: T5 (unstructured; N), T6 (hairpin; HP), and T7 (G-quadruplex; G4). Reactions were incubated at 35°C at specified time points.

In addition, we analyzed the effect of two RPA derivatives, RPAcore and RPA^ΔW^, on the Polα_CD_ activity on the normal and HP templates. Both truncations severely affected RPA’s ability to stimulate Polα_CD_ during HP bypass (Fig. 3B, lanes 3, 4, 7, and 8, compared to Fig. 3A, lanes 8 and 9). These results indicate that in the primosome, Polα makes the most important contacts with RPA.

### RPA does not assist human Polε_CD_ during DNA synthesis across a hairpin

For comparison, we analyzed the effect of RPA on activity of the catalytic domain of human DNA polymerase ε (Polε_CD_) under the same conditions as for Polα_CD_. Polε is specialized in synthesizing the leading strand, has a potent 3’→5’ exonuclease activity, and works in the complex with a CMG helicase (45–47). As for Polα_CD_, HP and G4 structures present a strong obstacle for Polε_CD_ (Fig. 3C). The pausing of DNA synthesis on HP and G4 templates was previously reported for yeast Polε (48) and human Polε_CD_ (49), respectively.

Strikingly, RPA does not help Polε_CD_ to bypass HP and shows an inhibitory effect on both normal and structured templates (Fig. 3C). Clearly, RPA can unfold HP ahead of Polε, as was observed in reactions with Polα, and that Polε_CD_ can displace RPA (Fig. 3C, lanes 1-4). However, RPA does not help Polε to bypass the HP motif. This fact points to the importance of cooperation between a DNA polymerase and RPA upon template handover between them. Of note, HP and especially G4 increase the level of products with a length below 16 nucleotides (Fig. 3C, lanes 5, 6, 9, and 10, in comparison to lanes 1 and 2), which points to the shift in the balance of polymerase/exonuclease activities toward the latter.

These results imply a different mode of RPA interaction with Polε versus Polα or to the absence of direct interaction. RPA might be of little importance for Polε because it receives the parental DNA strand directly from a CMG helicase.

### Structural studies revealed a dynamic complex between RPA and Polα connected by DNA

We prepared the complex for cryo-EM studies containing stoichiometric amounts of RPAcore^ΔP^, Polα_CD_, and the template:primer T9:P4 with a 24 nucleotides long 5’-flap (*SI Appendix*, Table S1). To increase the stability of the Polα complex with T:P, we used a chimeric RNA-DNA primer with a dideoxy-cytidine at the 3’-end to block DNA synthesis (37). Thus, the complex represents the early steps of DNA synthesis when Polα adds the third deoxynucleotide to the RNA primer generated by primase.

During data processing, it was found that most particles have a poor density for RPAcore except the W-domain, firmly attached to Polα (*SI Appendix*, Fig. S3). Such particles were filtered from further processing using different approaches, such as 2D and 3D classifications, *ab initio* 3D reconstruction, and heterogeneous refinement. Local refinement significantly improved the quality of the RPAcore^ΔP,W^/DNA map, resulting in the reconstruction at a resolution of 4.21 Å. For Polα_CD_/W/DNA, local refinement gave the reconstruction at a resolution of 3.49 Å, where Polα_CD_ shows a similar conformation as in the cryo-EM structure of the primosome elongation complex (10). It is worth mentioning that the entire complex side view projection ensembles a fish-like shape where RPAcore^ΔP,W^ forms a flexible tail fin, which explains its fuzzy density in the global 3D refinements (*SI Appendix*, Fig. S3).

RPA32 mediates the interaction between RPAcore and Polα_CD_ (Fig. 4). The W-domain of RPA docks on the α-helix (residues 615-629) of the Polα exonuclease domain, making hydrophobic and hydrophilic contacts including five hydrogen bonds (Fig. 4B; Polα_CD_ domains are shown in *SI Appendix*, Fig. S4A). Similar binding sites for the W-domain defined by only an α-helix were reported for DNA repair proteins uracil-DNA glycosylase UNG2 and a DNA translocase SMARCAL1 (28, 29). Moreover, the wHTH2 domain of CST, ssDNA-binding complex CTC1/STN1/TEN1 involved in telomere maintenance, structurally resembles the W-domain of RPA and binds Polα at the same site (50). In addition, four residues (198–201) of the linker, connecting the D- and W-domains of RPA32, dock into a hydrophobic pocket on a β-barrel at the N-terminal domain of Polα (Fig. 4B). At the third point of contact, the β-turn coil between two β-strands of the D-domain protrudes into a depression on the Polα N-terminal domain (*SI Appendix*, Fig. S4B) and works as a pivot allowing RPA rotation relative to Polα (*SI Appendix*, Fig. S3C). Such conformational flexibility might play a role in DNA handover from RPA to Polα.

**Figure 4.**
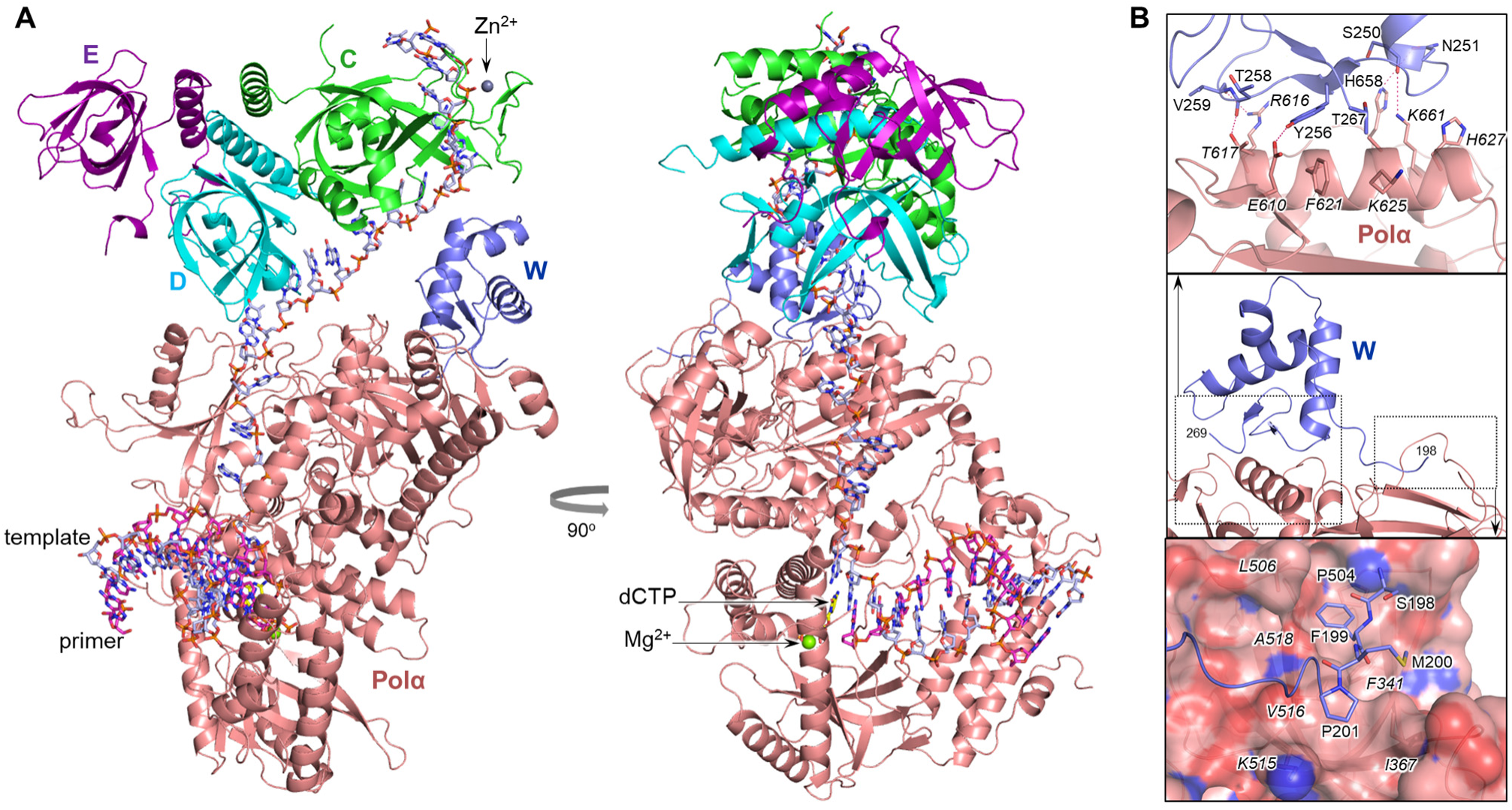
Structural basis for RPA-Polα cooperation during DNA synthesis. (**A**) Overall view of RPAcore/Polα_CD_/DNA complex. Polα and RPA domains C, D, E, and W are colored salmon, green, cyan, purple, and slate, respectively. The carbons of the DNA template, primer, and incoming dCTP are colored light blue, magenta, and yellow, respectively. (**B**) The close-up view of the W-Polα interaction interface. The residues involved in the interaction are shown as sticks; some main-chain atoms are not shown for clarity. Polα residues are labeled in italics. Dashed hot-pink lines depict H-bonds. Polα surface is represented at 20% transparency.

The structure shows a continuous engagement of the template DNA with RPAcore and the catalytic domain of Polα (Fig. 4A), which prevents the formation of secondary structures. A stretch of a DNA template between the Polα active site and RPA is in the cleft between the N-terminal and exonuclease domains of Polα (*SI Appendix*, Fig. S4). In addition, the disordered linker (amino acids 810-831) connecting the N-terminal and palm domains of Polα, is assumed to embrace this DNA region (*SI Appendix*, Fig. S4A). In general, human RPAcore interacts with DNA similarly as was described for RPA from *Saccharomyces cerevisiae* and the fungus *Ustilago maydis* (*SI Appendix*, Fig. S5) (31, 51), with a C-domain playing the main role in DNA binding.

### The W-domain stabilizes the RPAcore/DNA/Polα complex

We employed Bio-Layer Interferometry to determine the role of the W-domain in the interaction of RPA with DNA and Polα. With this approach, the rate constants of complex formation (*k*_on_), dissociation (*k*_off_), and the dissociation constant (*K_D_*) can be obtained. The intact RPA with all four DNA-binding domains is unsuitable for this assay due to very tight DNA binding, resulting in hours-long dissociation and notable ageing of solutions. We used RPA variants with a truncated N-terminal part of RPA70 to remove DNA-binding domains A and B. These domains are more dynamic on ssDNA with faster on-off kinetics than RPAcore (52). A pre-annealed T:P with a biotin at the 5′-tail of a template (*SI Appendix*, Table S1) was loaded on a streptavidin biosensor and dipped into solutions containing RPA variants at different concentrations.

First, we determined the affinity of RPAcore^ΔP,W^ to a 35-mer template annealed to an 11-mer primer (template makes a 24-mer 5’-tail) and obtained a *K_D_* value of 90 nM (Table 1). Next, we analyzed the effect of Polα on the RPAcore^ΔP,W^/DNA interaction. The DNA sensor was preloaded with Polα_CD_ in one well and then moved to the well containing RPAcore^ΔP,W^ in addition to the same concentration of Polα_CD_ so that the interaction of the RPAcore variant with DNA/Polα could be analyzed (*SI Appendix*, Fig. S6A). Strikingly, the *K_D_* value increased by almost 30-fold (up to 2.5 µM), mainly due to the more than 10-fold increase of a *k*_off_ value (Table 1). This result indicates a competition between Polα_CD_ and RPAcore^ΔP,W^ upon T:P binding.

**Table 1.** The effect of the W-domain and Polα_CD_ on interaction of RPAcore^ΔP^ with a template:primer.

| RPA variant | Pol$\alpha_{CD}$ | $k_{off} \times 10^{-3}$<br>sec$^{-1}$ | $k_{on}$<br>mM$^{-1}$ sec$^{-1}$ | $K_D^a$<br>nM |
| --- | --- | --- | --- | --- |
| RPAcore$^{\Delta P, W}$ | - | 24.6 ± 3.3 | 272 ± 22 | 90.2 ± 4.5 |
| | + | 266 ± 12.5 | 106 ± 9.5 | 2516 ± 108 |
| RPAcore$^{\Delta P}$ | - | 28.7 ± 3.2 | 232 ± 9.1 | 124 ± 15 |
| | + | 5.7 ± 0.35 | 115 ± 19 | 50.3 ± 6.6 |

<sup>a</sup> $K_D$ values are obtained by dividing $k_{off}$ by $k_{on}$ . Data are presented as mean ± SD (n = 3).

Compared to RPAcore^ΔP,W^, the variant with an intact W-domain (RPAcore^ΔP^) showed slightly lower affinity to T:P (*K_D_* = 124 nM, Table 1). This result indicates that the W-domain does not interact with DNA. Remarkably, in the presence of Polα_CD_, the affinity of this RPA variant to DNA increased 2.5-fold (*K_D_* = 50 nM), with a 5-fold reduction of the *k*_off_ value, which is inversely proportional to the complex stability (Table 1, binding kinetics are shown in *SI Appendix*, Fig. S6B). Thus, the W-domain causes a switch in the cooperativity between RPAcore and Polα, turning it from negative to positive and providing a ∼50-fold gain in the stability of the RPAcore/Polα/DNA complex. This complex may exist *in vivo* when AB-domains are displaced from DNA by the moving primosome (see Discussion). A stimulatory effect of human Polα on the RPAcore-DNA interaction agrees with studies conducted with yeast proteins showing that RPA stabilizes the Polα/DNA complex (53).

## DISCUSSION

Our studies uncovered a critical role of RPA in DNA synthesis across hairpins by primosome. The important factors required for RPA-assisted hairpin bypass by Polα are the high affinity of RPA to DNA and the interaction of the W-domain with the catalytic domain of Polα. This interaction dramatically increases cooperativity between RPAcore and Polα upon T:P binding and keeps RPA in close proximity to Polα, which allows for a seamless handover of unfolded DNA. RPA does not stimulate a DNA-polymerase activity of primosome on the unstructured template and rather facilitates synthesis termination at the RNA-DNA primer length of 32 nucleotides.

Current structural data allow the entire RPA/primosome/DNA elongation complex to be modeled, revealing its extended shape (*SI Appendix*, Fig. S7). In such a compact form, only 26 nucleotides of an unpaired DNA template are covered by RPA and Polα. Potentially, the linker between the B- and C-domains of RPA can extend resulting in a more significant footprint of RPA on DNA. The model is consistent with a mode of primosome integration into a replisome (8) and with the results of biochemical studies presented here, pointing to the main role of Polα in RPA-primosome interactions during DNA synthesis.

This work clarifies in what order RPA domains are displaced from a template as DNA synthesis progresses. According to structural data, RPAcore plays an essential role during primer synthesis by directing the unfolded DNA template into the active site of Polα. RPAcore binds to Polα/DNA with higher affinity than the AB-domains bind to DNA (*K_D_* values are 50 nM (Table 1) and 114 nM (54), respectively). Consistently, in the cryo-EM structure of yeast RPA, complexed with 100-mer ssDNA, only RPAcore was clearly visible in the complex with DNA, despite the absence of Polα (51). The decreased stability of the AB/DNA complex is probably due to the flexible connection of DNA-binding domains A and B, each having a one to two orders of magnitude lower affinity to DNA in comparison to their tandem (54). These data point to a scenario where the N-terminal part of RPA70 is displaced from ssDNA first, probably due to sequential dissociation of A- and B-domains. Of note, an upstream RPA molecule may have higher chances to be displaced from DNA than RPA attached to primosome.

## MATERIALS AND METHODS

### Oligonucleotides for functional studies

Sequences of all oligonucleotides are provided in *SI Appendix*, Table S1. Oligonucleotides were obtained from IDT Inc. A 12-mer chimeric RNA-DNA primer (P1) was obtained by ligation as described elsewhere (17).

### Protein expression and purification

Purification of the human primosome and the Polα catalytic core have been described elsewhere (44). As previously reported, the N-terminus of p180 (residues 1-334) has been deleted because it is poorly folded and is not required for activity (10, 50). Purification to homogeneity of the catalytic domain of human Polε (Polε_CD_, residues 28-1194) was conducted according to (55).

Genes encoding three subunits of human RPA were cloned into pASHSUL-1 plasmid (56), resulting in a poly-cistronic expression vector with the following order of subunits: RPA70, RPA32, and RPA14. A cleavable His-Sumo-tag was placed at the N-terminus of RPA70. RPA truncation variants were obtained according to (57). Human RPA and its variants were expressed in *Escherichia coli* strain BL21 (DE3) at 16 °C for 15 hours following induction with 0.3 µg/ml anhydrotetracyclin. Afterward, cells were harvested by centrifugation at 4,000 g for 15 min, aliquoted, and kept at -80 °C.

All RPA variants were purified according to the same protocol, including chromatography on a Ni-IDA column (Bio-Rad), His-Sumo tag digestion during overnight incubation on ice, and two additional chromatographic steps using Heparin HP HiTrap and monoQ 5/50 GL columns (Cytiva). Finally, samples were concentrated and flash-frozen in aliquots. The purity of samples was analyzed by SDS-PAGE (*SI Appendix*, Fig. S1). Protein concentrations were estimated by measuring the absorbance at 280 nm and using the extinction coefficients calculated with ProtParam (58).

### Primosome activity assay

DNA-synthetic activity of primosome was tested in a 10 μl reaction containing 0.2 μM T:P, 0.2 µM enzyme, 1µM RPA or its variants, 10 μM dNTPs, 0.1 µM [α-^32^P]-dCTP (3000 Ci/mmol; PerkinElmer, Inc.), and the buffer: 30 mM Tris-Hepes, pH 7.8, 120 mM KCl, 30 mM NaCl, 1% glycerol, 1.5 mM TCEP, 5 mM MgCl_2_, and 0.2 mg/ml BSA. The proteins were pre-incubated with T:P for 1 min on ice and for 10 sec at 35 °C, then the reaction was initiated by adding an equal volume of buffer solution containing dNTPs. Reactions were incubated in PCR tubes on a water bath for the indicated time points at 35 °C and stopped by mixing with an equal volume of formamide loading buffer (90% v/v formamide, 50 mM EDTA, pH 8, 0.02% Bromophenol blue), heated at 95 °C for 1 min, and resolved by 20% Urea-PAGE (UreaGel System (19:1 Acrylamide/Bisacrylamide), National Diagnostics) for 2.5 h at 3000 V. The reaction products were visualized by phosphorimaging (Typhoon FLA 9500, Cytiva). Experiments were repeated three times.

### DNA polymerase activity assay

DNA-synthetic activity of Polα_CD_ and Polε_CD_ was tested in a 10 μl reaction containing 0.25 μM T:P, 20 nM enzyme, 1µM RPA or its variants, 50 μM dNTPs, and the buffer: 30 mM Tris-Hepes, pH 7.8, 120 mM KCl, 30 mM NaCl, 1% glycerol, 1.5 mM TCEP, 5 mM MgCl_2_, and 0.2 mg/ml BSA. A 16-mer DNA primer contains a Cy5 fluorophore attached to the 5’-end. RPA was pre-incubated with T:P and dNTPs for 1 min on ice and for 10 sec at 35 °C, then the reaction was initiated by adding an equal volume of buffer solution containing enzyme. Reactions were incubated in PCR tubes on a water bath for the indicated time points at 35 °C and stopped by mixing with an equal volume of formamide loading buffer (90% v/v formamide, 50 mM EDTA, pH 8, 0.02% Bromophenol blue), heated at 95 °C for 1 min, and resolved by 20% Urea-PAGE (UreaGel System (19:1 Acrylamide/Bisacrylamide), National Diagnostics). The reaction products were visualized by imaging on Typhoon FLA 9500 (Cytiva). Experiments were repeated three times.

### Binding studies

The binding studies were conducted at 23 °C using Octet K2 (Sartorius AG), as previously described (59). This device uses Bio-Layer Interferometry technology to monitor molecular interactions in real-time. The 35-mer template with a biotin-TEG at the 5’-overhang was annealed to an 11-mer primer (*SI Appendix*, Table S1) and immobilized on a streptavidin-coated biosensor (SAX, Sartorius AG). A chimeric RNA-DNA primer demonstrating the highest affinity to Polα (37) was utilized to saturate DNA sensors with Polα_CD_ at a minimal concentration. Dideoxy-cytidine was placed at the primer 3′-end to prevent DNA polymerase reaction. SAX sensors were loaded with oligonucleotide-biotin at 50 nM concentration for 7 min at 500 rpm; then, sensors were blocked by incubating with 10 µg/ml biocytin for 2 min.

Binding studies were conducted in a 96-well microplate (Greiner Bio-One) in buffer containing 25 mM Tris-Hepes, pH 7.8, 120 mM KCl, 30mM NaCl, 2.5 mM MgCl_2_, 50 µM dTTP, 2 mM TCEP, and 0.002% Tween-20. The first six wells in a row contained only the buffer, and the subsequent six wells contained two-fold dilutions of the RPA variant in the same buffer. Before and after incubation with RPA (association step), the DNA-sensor was placed into the same RPA-free well (distinct for each RPA concentration), resulting in baseline and dissociation steps, respectively.

Upon binding studies in the presence of Polα_CD_, the DNA sensor was moved first to the well containing 100 nM Polα_CD_. The next well, where the association step was carried out, contained an RPA variant as well as 100 nM Polα_CD_ to maintain an equilibrium between sensor-bound and free Polα_CD_. To observe the dissociation of an RPA variant from the Polα/DNA complex, a sensor was moved to the same well where it was pre-loaded with Polα_CD_. RPAcore and its variant containing a W-domain were used at concentrations of 3.6 µM and 200 nM, respectively, exceeding the corresponding *K_D_* values. Data Analysis HT software (ver. 11.1, Sartorius AG) was used to calculate binding constants (*k*_on_, *k*_off_, and *K_D_*). The average value and standard deviation were calculated from three independent experiments.

### Cryo-EM sample preparation, data collection, and processing

The complex RPAcore^ΔP^/Polα_CD_/T9:P4 was prepared by mixing stoichiometric amounts of pure proteins and a template:primer in a buffer containing 20 mM Hepes-Na, pH 7.5, 100 mM KCl, 20 mM NaCl, 1% glycerol, 0.5 mM dCTP, 0.9 mM MgCl_2_, 0.2 mM CaCl_2_, 1 mM TCEP. The holey carbon copper support electron microscopy grids (C-Flat 1.2/1.3, 300 mesh) were glow-discharged for 60 s at 15 mA negative current (PELCO easiGlow) before use. CHAPSO was added to the complex at the final concentration of 3.3 mM before depositing a 3 µl sample at the protein concentration of 4 mg/ml or 2.5 mg/ml onto the glow-discharged grid. The grids were blotted for 3 s at 4°C with a bloat force of 5 and 95% relative humidity and plunged frozen into liquid ethane using a Vitrobot Mark IV (Thermo Fisher Scientific). A total of 18,531 raw movies were acquired using a 200 keV Glacios fringe-free corrected and energy-filtered cryo-transmission electron microscope equipped with a Falcon 4i direct electron detector using the EPU automated acquisition software (Thermo Fisher Scientific) with ‘‘Faster Acquisition’’ mode (AFIS) enabled. A slit width of 10 eV was used for the Selectrics energy filter, in addition to a 20 µm condenser aperture allowing the collection of three shots per hole. Data were collected at an effective pixel size of 0.72 Å/pixel, a defocus range from -0.8 to -2.4 µm, and an exposure time of 4.454 s with a total dose of 60.0 e /Å^2^.

The data processing pipeline outlined here is schematized in *SI Appendix*, Fig. S3. Data were processed using CryoSPARC (ver. 4.6) (60). The movies were aligned via patch motion correction, and their CTF parameters were estimated using the patch CTF estimation. 972,867 particles were extracted from curated micrographs with a box size of 450 Å using the blob- and template-based particle-picking approaches. Three rounds of 2D classification were carried out, resulting in 313,894 selected particles. Several rounds of *ab initio* 3D reconstruction and heterogeneous refinement were conducted before and after the local CTF refinement to clean the dataset from the low-quality particles, ending with 105,167 particles. The poor quality of the cryo-EM density map for RPAcore/DNA pointed to its flexibility. 3D classification revealed four main conformers due to RPAcore^ΔP,W^ rotation and tilting relative to Polα (*SI Appendix*, Fig. S3C). The class with the highest quality of density map for RPAcore/DNA was selected for further processing. At this step, it was clear that the W-domain of RPA is firmly attached to Polα.

Soft masks for Polα_CD_/W/DNA, RPAcore^ΔP,W^/DNA, and DNA were created in UCSF ChimeraX (ver. 1.7.1) (61) and used for particle subtraction, local refinement, and 3D classification. 24,611 particles were obtained after additional cleaning by several rounds of *ab initio* 3D reconstruction and heterogeneous refinement and used for 3D classification with a mask applied to a DNA stretch located between the C- and D-domains of RPA. The classes with continuous density for DNA were selected resulting in 18,529 particles with an overall cryo-EM density map resolution of 3.5 Å (*SI Appendix*, Fig. S3C). Using the density-subtracted particles for Polα_CD_/W/DNA, local refinement resulted in the reconstruction at 4.21 Å for RPAcore^ΔP,W^/DNA. Using the density-subtracted particles for RPAcore^ΔP,W^/DNA, the local refinement generated a reconstruction for Polα_CD_/W/DNA at 3.49 Å. Local maps were combined in Phenix (ver. 1.21.2) (62) using the consensus map as a guide.

### Model building and refinement

The initial model was built by the rigid-body fitting of the three starting models, including Polα_CD_ with duplex (4QCL), RPAcore (1L1O) and the W-domain generated with AlphaFold 3 (63), into the final overall map using the ChimeraX. All further manual adjustments of the initial model, adding the single-strand part of the DNA template and the amino-acid residues missed in the initial model, were performed using the Coot (64). Poor quality of EM density for Polα regions 810-831 and 883-895 and RPA32 regions 35-42 and 179-197 precluded unambiguous placement of corresponding residues. After several rounds of manual model building, the structure was refined using the “real-space refinement” module of Phenix. Refinement and validation statistics are provided in *SI Appendix*, Table S2. PyMol Molecular Graphics System (version 1.8, Schrödinger, LLC) was used for figure preparation.

## Data sharing statement

The RPAcore/Polα_CD_/DNA complex structure and the corresponding map are deposited in the Protein Data Bank under accession codes 9MJ5 and EMD-48312, respectively.

## Acknowledgements

This work was supported by the National Institute of General Medical Sciences grant R35GM152032 to T.H.T. The UNL Cryo-EM core facility (RRID:SCR_025012) is supported by Federal funding from NIH COBREs (NCIBC) grant GM113126. We thank J. Lovelace for providing resources for model refinement and K. Jordan for editing this manuscript.

## Author contributions

A.G.B., L.M.M. and N.D.B. conceived experiments and analyzed the data.

A.G.B. and E.R.R. prepared the cryo-EM grids, collected the data, and reconstructed the cryo-EM maps.

T.H.T. built the model and supervised the project. A.G.B. wrote the manuscript, with contributions and critical comments from the other authors.

