## Supplemental information for "Human primosome requires replication protein A when copying DNA with inverted repeats"

for the article

**This file includes:**

Tables S1 and S2
Figures S1 - S7

**Table S1. Oligonucleotides used in this study**

| Name | Sequence [a] | Application | Length |
|---|---|---|---|
| T1 | ATTTTGAAGAGAAATACTTTAAATCAGTCTGGAATGATGAAGATTACTAGTGAAGATTCTGAGCGTCTTAATCTAAGTACAGGTCGTGCCGCCAAAAA[b] | Primer extension | 98 |
| T2 | ATTTTGAAGAGAAATACTTTAAATCAAATGATGAAGATATCTAATCTATGGTCGCTCCATTCAGGAGCGACCTGTAGTACAGGTCGTGCCGCCAAAAA[c] | | 98 |
| T3 | ATTTTGAAGAGAAATACTTTAAATCAAATGATGAAGATATAGCGTCTAATCTATGAGGGTGGGTAGGGTGGGTGTAGTACAGGTCGTGCCGCCAAAAA | | 98 |
| T4 | CTGAGCGTCTTAATCTAAGCACAGGTCGTGCCGCCAAAAA | | 40 |
| P1 | ppp*GGCGGCACG*ACC[d] | | 12 |
| T5 | GTCTGGAATGATGAAGATTACTAGTGAAGATTCTGAGCGTCTTAATCTAAGCACTCGCTATGTTTTCAAGTTT | | 73 |
| T6 | AATGATGAAGATATCTGGTCGCTCCATTCTGGAGCGACCTCTTAATCTAAGCACTCGCTATGTTTTCAAGTTT | | 73 |
| T7 | GTCTGGAATGATGATGAGGGTGGGTAGGGTGGGTGAGCGTCTTAATCTAAGCACTCGCTATGTTTTCAAGTTT | | 73 |
| P2 | /Cy5/CTTGAAAACATAGCGA | | 16 |
| T8 | /Biotin/AATCTAGTAACATAGTATACATAAGCGCTCCAGGC | Binding kinetics | 35 |
| P3 | *GCCUGGAG*CG/3ddC/[e] | | 11 |
| T9 | AATCTAGTAACATAGTATACATAGGCGCTCCAGGC | Cryo-EM | 35 |
| P4 | *GCCUGGAGC*G/3ddC/ | | 11 |

[a] Sequences are listed in order from 5'-end to 3'-end.
[b] The regions complementary to a primer are underlined.
[c] The structured regions of a DNA template are in red (T2 and T6 form a hairpin, T3 and T7 form a G-quadruplex).
[d] The ribonucleotides are in italics; ppp indicates the 5'-triphosphate group.
[e] 3'-dideoxy-cytidine.

**Table S2. Cryo-EM data collection, refinement and validation statistics for the RPAcore/Polα$_{CD}$/DNA complex.**

| | |
|---|---|
| **Data collection and processing** | |
| Microscope | Glacios |
| Voltage (keV) | 200 |
| Detector | Falcon 4i |
| Magnification | 165,000 |
| Electron dose ($e^-$/Å$^2$) | 60 |
| Pixel size (Å) | 0.72 |
| Defocus range (µm) | -0.8 to -2.4 |
| Symmetry imposed | C1 |
| Resolution (Å) | 3.5 |
| FSC threshold | 0.143 |
| **Refinement** | |
| Initial model used (PDB code) | 4QCL, 1L1O |
| Non-hydrogen atoms | 11919 |
| Protein residues | 1378 |
| RNA/DNA nucleotides | 42 |
| Cofactors/ions | 3 |
| R.m.s. deviations | |
| Bond lengths (Å) | 0.003 |
| Bond angles (°) | 0.61 |
| **Validation** | |
| MolProbity score | 1.6 |
| Clash score | 5.9 |
| Poor rotamers (%) | 0.65 |
| Ramachandran plot (%) | |
| Favored | 96.11 |
| Allowed | 3.81 |
| Disallowed | 0.07 |
| Fit to map ($CC_{mask}$) | 0.83 |
| **Accession codes** | 9MJ5, EMD-48312 |

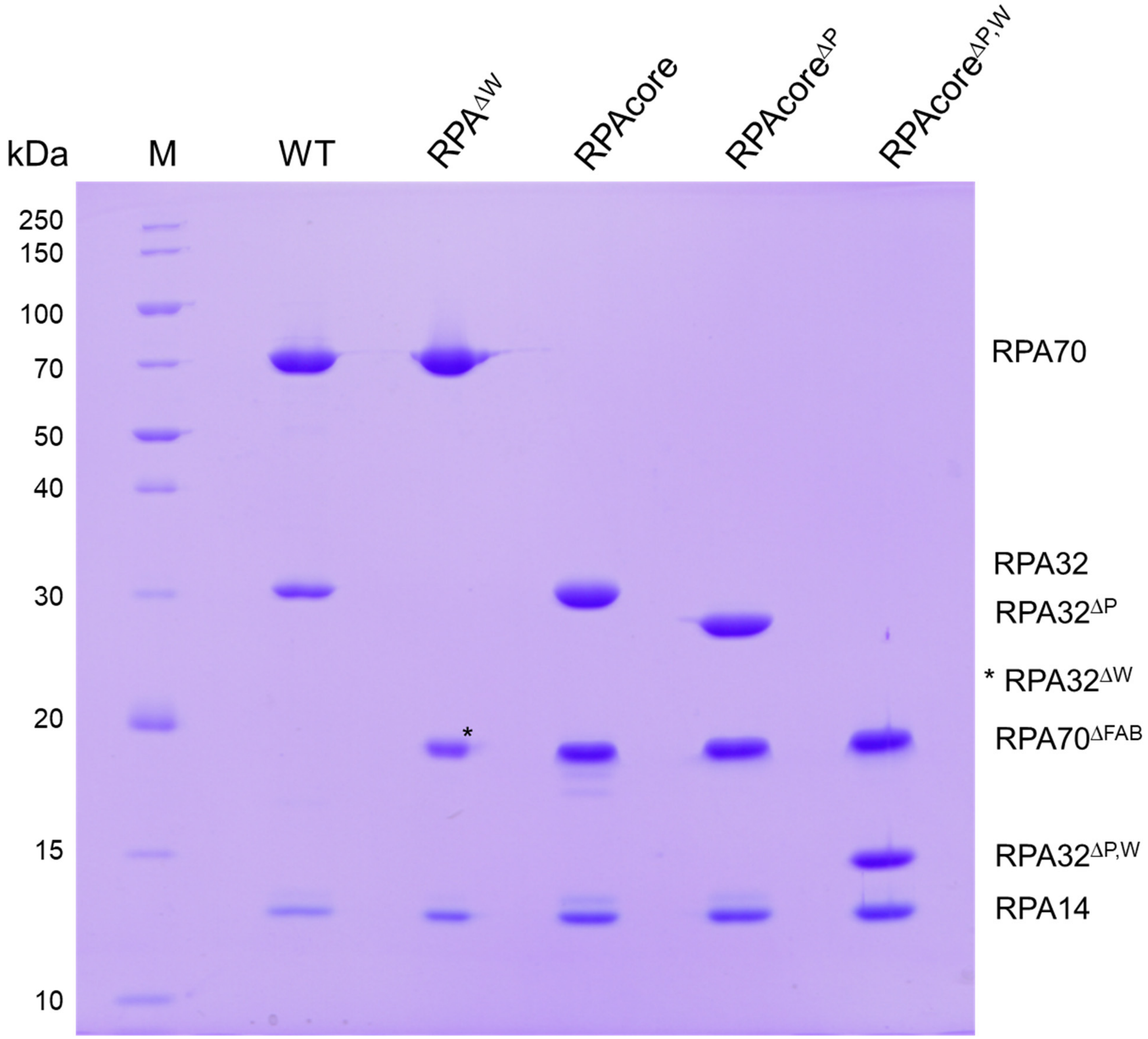


**Figure S1. Analysis of purity of human RPA and its variants.** Proteins were separated by 13% SDS-PAGE and stained by Coomassie Brilliant Blue R-250.

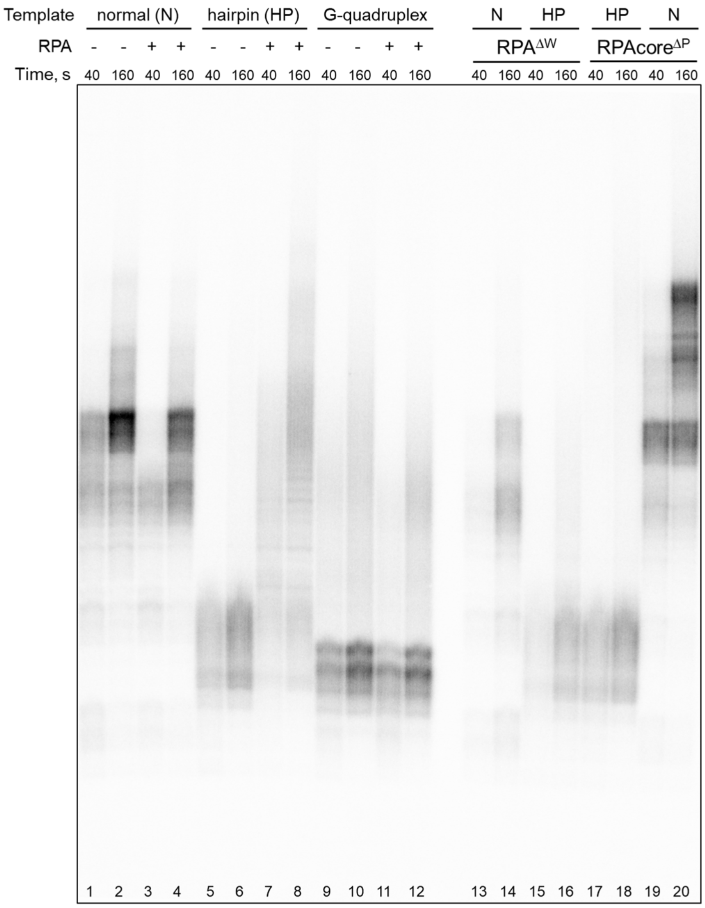


**Figure S2. Analysis of selected primosome reactions on one gel.** A deletion of the W-domain has an inhibitory effect on DNA synthesis by the RPA-primosome complex on the unstructured template and in the presence of a hairpin (lanes 3-6, in comparison to lanes 13-16). A deletion of the RPA70 N-terminus (contains F-, A-, and B-domains) has a stimulatory effect on DNA synthesis on the unstructured template (lanes 3 and 4, in comparison to lanes 19-20) and an inhibitory effect on the template containing a hairpin (lanes 5 and 6, in comparison to lanes 17-18). The 98-mer DNA templates of different structure were annealed to a 12-mer chimeric RNA-DNA primer (P1) containing the triphosphate at the 5'-end. The following templates were used: T1 (unstructured or normal), T2 (hairpin), and T3 (G-quadruplex). Reactions were incubated at 35ºC at specified time points.

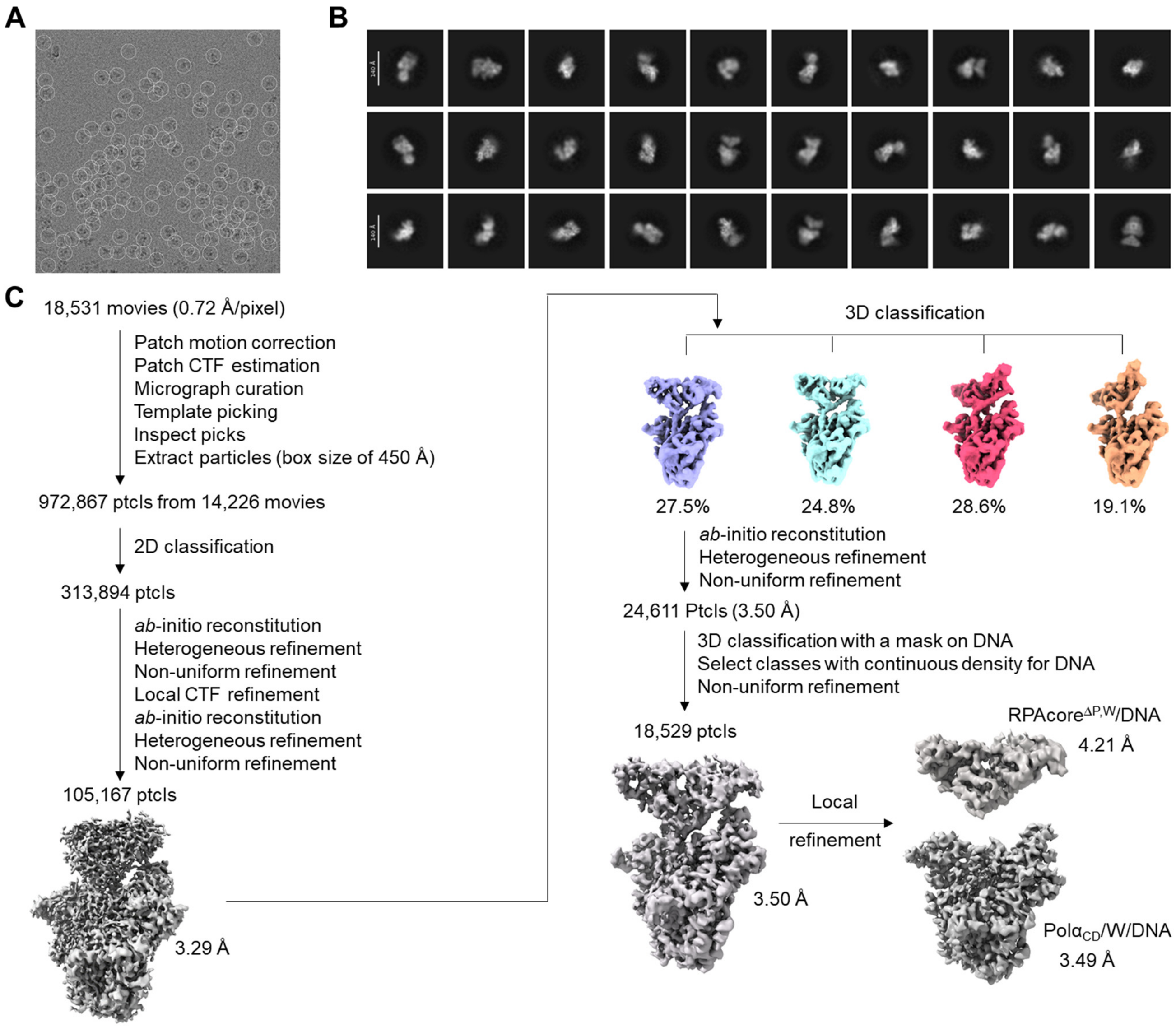


**Figure S3. Cryo-EM processing pipeline for the RPAcore/Polα$_{CD}$/DNA complex. (A)** Representative micrograph with particles (ptcls) passed inspection before extraction. (**B)** 2D classification averages selected after the third round. (**C)** Cryo-EM processing pipeline.

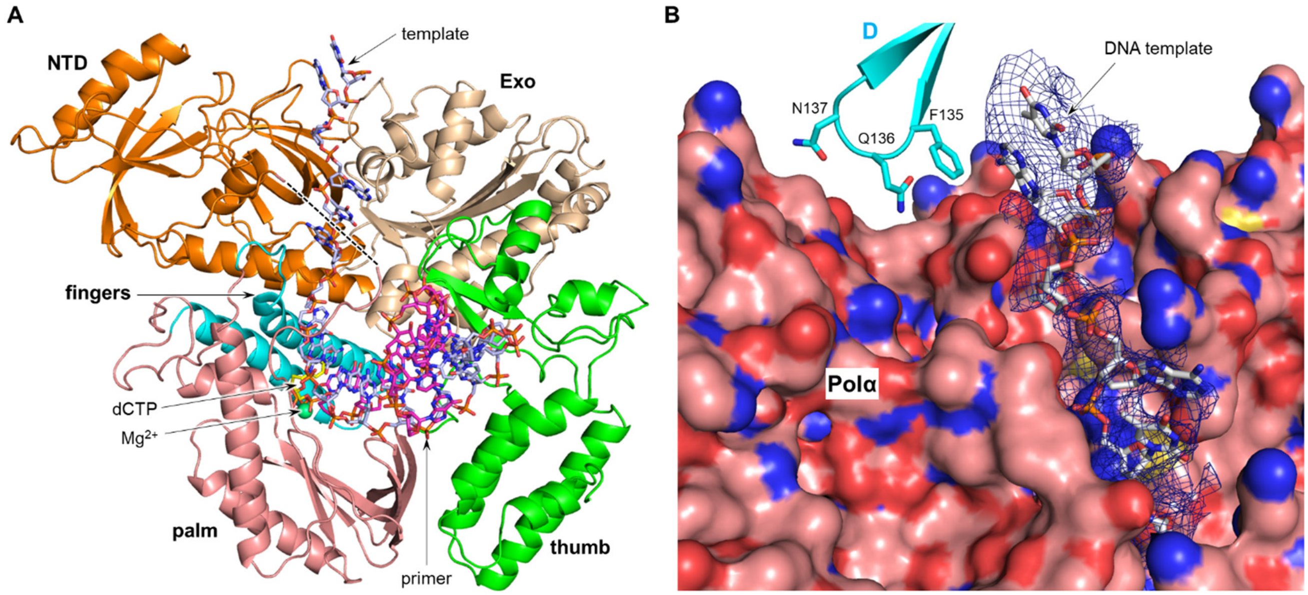


**Figure S4. The cleft between the N-terminal and exonuclease domains of Polα directs a DNA template from RPA to the Polα active site.** **(A)** The overall view of Polα/DNA/dCTP complex. Polα domains N-terminal (NTD), exonuclease (Exo), fingers, palm, and thumb are represented as cartoon and colored orange, wheat, cyan, salmon, and green, respectively. Template, primer, and dCTP are shown as sticks with carbons colored light-blue, magenta, and yellow, respectively. The disordered linker (residues 810-831) connecting the N-terminal and palm domains is represented by dashed line. RPA and DNA in complex with it are not shown for clarity. (**B**) The close-up view of a DNA template in the Polα cleft and the Polα-RPA contact area formed by NTD and the D-domain. Polα and DNA are represented as surface and sticks, respectively. Three RPA residues in proximity to Polα NTD are shown as sticks. Atoms of nitrogen, oxygen, sulfur, and phosphor are colored blue, red, yellow, and orange, respectively. Carbons of protein and DNA are colored salmon and light-blue, respectively. Electronic density of DNA is shown as a blue mesh.

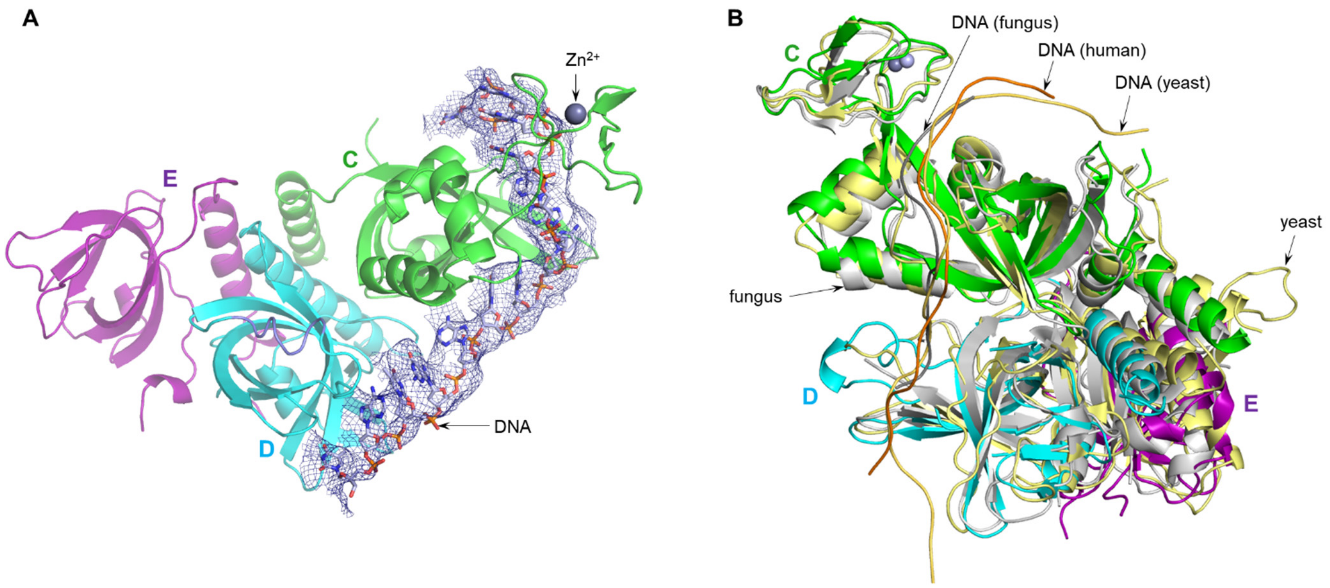


**Figure S5. Human RPAcore binds DNA similar to other species. (A)** DNA interacts with the C- and D- domains of human RPAcore. DNA and its electronic density are shown as sticks and blue mesh, respectively. Protein is represented as cartoon at 20% transparency. **(B)** Comparison of a DNA path on RPAcore from different species. The human RPAcore/DNA complex was aligned with corresponding complexes, containing yeast (pdb ID 6I52) and fungal RPA (pdb ID 4GNX), with rmsd of 2.3 Å and 2.6 Å, respectively. DNA is represented as ribbon. The C, D, and E domains of human RPAcore are colored green, cyan, and purple, respectively; DNA is colored orange. The yeast and fungal RPAcore molecules (and DNA in the complex with them) are colored pale-yellow and gray, respectively. $Zn^{2+}$ ions in human and fungal RPA are shown as spheres and colored light-blue and gray, respectively. For clarity, the AB-domains of fungal RPA as well as DNA in complex with them are not shown.

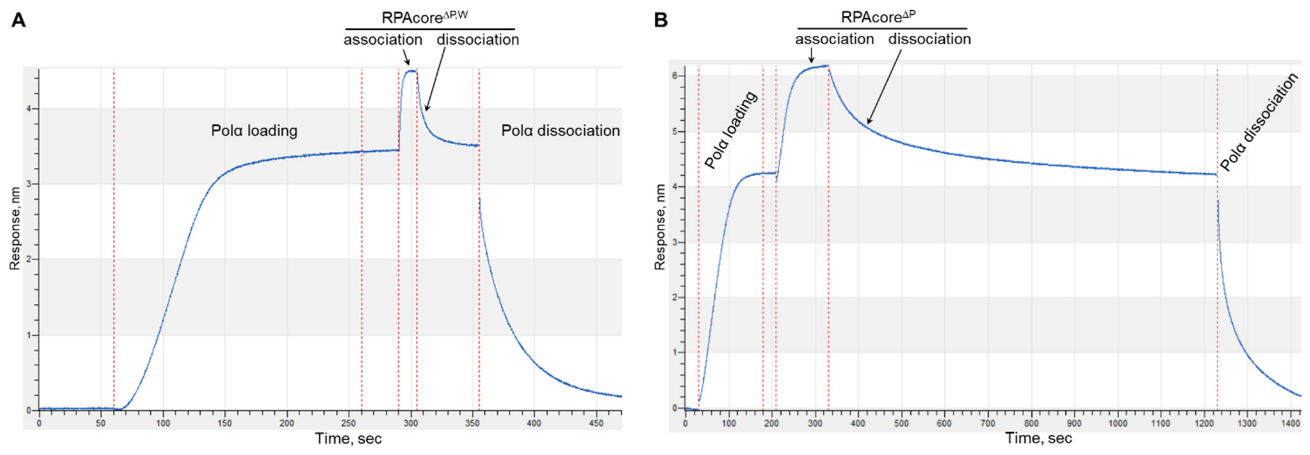


**Figure S6. DNA-binding studies of RPAcore$^{\Delta P,W}$ (A) and RPAcore$^{\Delta P}$ (B) in the presence of Polα$_{CD}$.** RPAcore$^{\Delta P}$ and its variant missing the W-domain dissociate from the DNA/Polα sensor in 15 min and 50 sec, respectively.

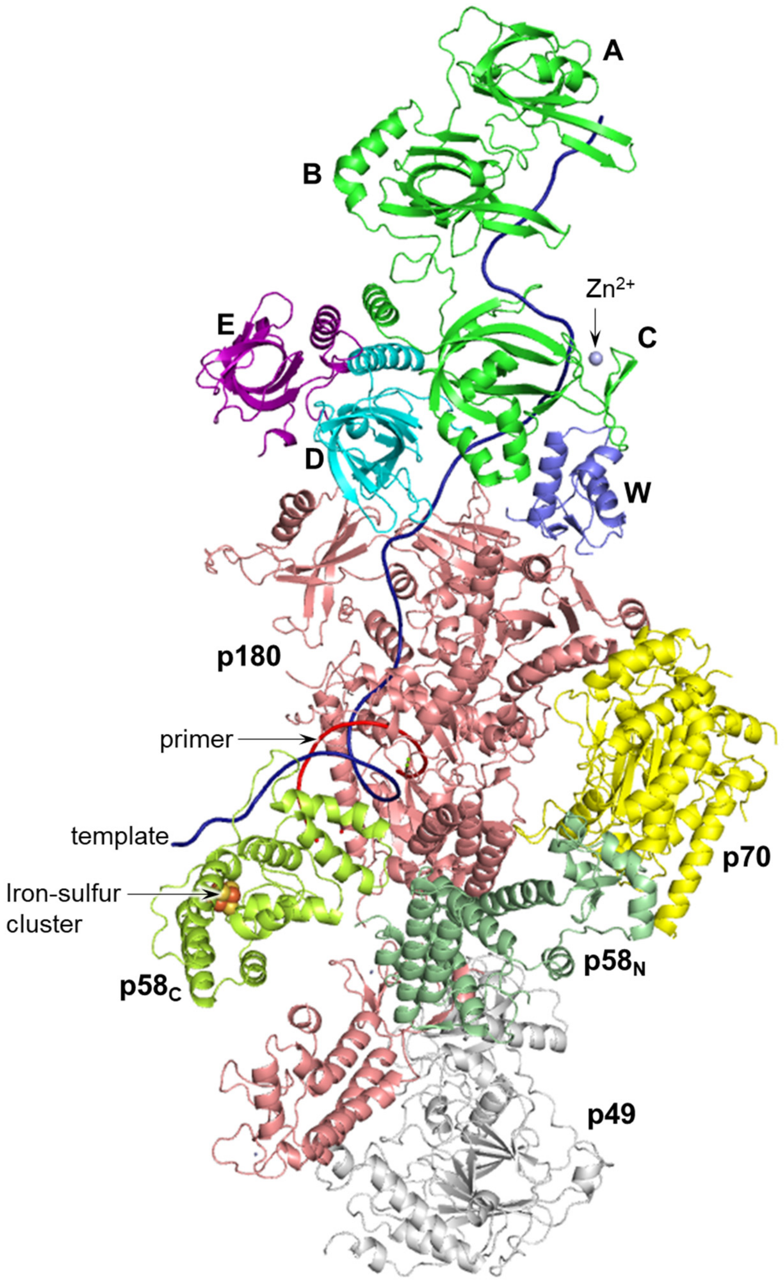


**Figure S7. The model of RPA/primosome elongation complex.** The complex RPAcore/Polα$_{CD}$/DNA was superimposed onto the primosome elongation complex (pdb ID 8D9D) using Polα$_{CD}$ for alignment (rmsd of 0.56 Å). The model for human RPA in complex with DNA was generated using AlphaFold 3 Server and superimposed onto the modeled complex RPAcore/primosome/DNA using RPAcore for alignment (rmsd of 0.94 Å). From the RPA/DNA model generated by AlphaFold, only the AB-domains and 11-mer DNA in complex with them are shown. The F-domain of RPA is very flexible and is not shown for clarity. Human Polα comprises subunits p180 (catalytic) and p70 (accessory); human primase comprises subunits p49 (catalytic) and p58 (accessory), which has N-terminal ($p58_N$) and C-terminal ($p58_C$) domains. $p58_C$ interacts with the primer 5'-end and regulates all steps of RNA-DNA primer synthesis.
